# Spontaneous Self-assembly of Amyloid β (1-40) into Dimers

**DOI:** 10.1101/659300

**Authors:** Mohtadin Hashemi, Yuliang Zhang, Zhengjian Lv, Yuri L. Lyubchenko

## Abstract

The self-assembly and fibrillation of amyloid β (Aβ) proteins is the neuropathological hallmark of Alzheimer’s disease. However, the molecular mechanism of how disordered monomers assemble into aggregates remains largely unknown. In this work, we characterize the assembly of Aβ (1-40) monomers into dimers using long-time molecular dynamics simulations. Upon interaction, the monomers undergo conformational transitions, accompanied by change of the structure, leading to the formation of a stable dimer. The dimers are primarily stabilized by interactions in the N-terminal region (residues 5-12), in the central hydrophobic region (residues 16-23), and in the C-terminal region (residues 30-40); with inter-peptide interactions focused around the N- and C- termini. The dimers do not contain long β-strands that are usually found in fibrils.

## Introduction

The self-assembly of amyloidogenic proteins is related to several neurodegenerative diseases(1–3). According to the amyloid cascade hypothesis, self-assembly of amyloid β (Aβ) is the primary model for the development of Alzheimer’s disease (AD) (1, 4). The final products of the amyloid self-assembly process are fibrillar structures that contain long β-strands(5–7), whereas Aβ monomers are largely unstructured(8–10), which leads to the question of how the conformational transition occurs during self-assembly.

Recent compelling evidence show that amyloid oligomers rather than fibrils are the most neurotoxic species (11–15). The neurotoxicity of Aβ oligomeric species have been attributed to intracellular, membrane, and receptor-mediated mechanisms (16–27). Various morphologies have been ascribed to oligomers, from spherical aggregates to filamentous (28, 29). It is proposed that oligomers form the critical entities, called nuclei, needed to transition to proto-fibril states before finally fibrillating (30). Spectroscopic characterization of Aβ oligomers revealed that they are composed of random coil secondary structure, which is able to transition to β-structure as the aggregation progresses (30–33). Sarkar *et al.* showed that the oligomer chemical shifts are very different from fibrils, in particular the N-terminal and the central segment (residues 22-29) (32). These finding are in line with the data from Ahmed *et al.*, which show that oligomers have disordered molecular conformations (30).

There are two principle alloforms of amyloid β proteins, Aβ (1-40) and Aβ (1-42), defined by the number of residues; with the former being the most abundant and the latter the most aggregation prone and neurotoxic (34–40). Despite the small structural difference (two amino acids) between the Aβ40 and Aβ42 alloforms, they display distinct behavior, although the structural basis for this is unknown (37–41). Hence, a detailed characterization of these oligomeric forms of Aβ is important for understanding neurotoxicity and pathology in AD. Recent studies have demonstrated that single-molecule approaches are powerful methods to study oligomers (42–45). Single-molecule techniques, such as AFM (46–51), tethered approach for probing inter-molecular interactions (TAPIN) (52, 53), and FRET (32), have shown that the early stage oligomers exhibit prolonged lifetimes and stabilities. Novel features of the interaction and self-assembly of Aβ40 and Aβ42 peptides were determined using single-molecule AFM-based force spectroscopy (47). However, due to their transient nature and heterogeneity many questions about the oligomer formation process structure and the structure and dynamics of Aβ oligomers are left unanswered (54, 55).

Computational simulations have been utilized to supplement the novel single-molecule techniques used to probe early stages of aggregation and, in some cases, elucidate the dynamics and mechanism of aggregation (50, 56–60). Computational studies of the dynamics of Aβ42 lead to the discovery that, in an aqueous environment, the protein mainly assumes α-helical structure (61). However, the helices are not stable and transition between structured and unstructured conformations multiple times. Further studies showed that Aβ42 is more structured compared to Aβ40 and has a less flexible C-terminal segment (57). These findings are in line with the comparison of Aβ40 and Aβ42 by Yang and Teplow, which showed that Aβ42 forms more stable conformations that tend towards β-structure and stable C-terminus (62). More recent simulations have revealed that the size and distribution of the early aggregates for Aβ40 and Aβ42 vary, the most common oligomer being dimers for the former and pentamers for the latter (63, 64). These results qualitatively reproduce the main features of oligomer size distributions measured experimentally (41, 65). Furthermore, Aβ42 displayed turn and β-hairpin structures that are absent in Aβ40.

Biased simulation strategies using a coarse-grained force field has also been employed to investigate the aggregation pathway (66). Zheng *et al.* demonstrate that the while pre-fibrillar oligomers typically consist of antiparallel β-structure they are distinct from fibrillar structures and very dynamic. These structural characteristics are also demonstrated for the Aβ40 dimer in the findings of Tarus *et al.*, which show that dimers are compact conformations with inter-peptide antiparallel β-structures (67). Similar observations were also reported by Watts *et al.* using a different force field (68). However, how the structures of oligomers contribute to neurotoxicity remain unclear. Leaving the fundamental questions related to the mechanism of oligomer self-assembly and dynamics unanswered. Which, in turn, has impeded the progress in the development of treatment for these diseases.

We recently characterized the conformational changes in monomers of Aβ42 peptide upon dimer formation using long time-scale all-atom molecular dynamics (MD) simulations (69). The simulations revealed that the dimer is very dynamic and resulted in a multitude of different conformations being identified. By utilizing the recently developed Monte Carlo pulling (MCP) approach (58), we were able to identify the most likely native conformations of the Aβ42 dimer, which generated statistically similar dissociation forces and interaction profiles as was observed in AFM experiments.

Here, we applied the developed MD simulation strategy to analyze the dimer formation of full-length Aβ40 protein using the specialized Anton supercomputer (70, 71). A variety of dimer conformations were identified, all with small segments of ordered structures and lacking the characteristic β-sheet structures found in amyloid fibrils. These dimers structures were then validated using MCP simulations and by comparing with stability and interaction data obtained from AFM-based force spectroscopy experiments. The validated dimer conformations were then used to compare Aβ40 and Aβ42 dimers and characterize the differences between the interaction of monomers and the resulting dimers.

## Materials and methods

### Monomer simulation procedure

To generate the initial structure of the monomers used for the dimer simulation, we conducted all-atom MD simulations using GROMACS ver. 4.5.5 (72) employing Amber ff99SB-ILDN force field(73) and the TIP3P water model(74). The initial monomer structures were adopted from NMR data(8) (PDB ID: 1AML) and an extra N-terminal Cys residue was added to mimic experimental sequence (69). The monomer was then solvated, neutralized with NaCl ions, and kept at 150 mM NaCl concentration. After which 500 ns NPT MD simulation, at 1 bar and 300 K, was carried out. Cluster analysis was then performed using the GROMOS method of clustering and root-mean square deviation (RMSD) as input for the protein backbone with a 3Å cut-off value, as previously described (50).

### Dimer simulation on the specialized supercomputer Anton

The initial dimer conformations were prepared in the Maestro software package (Schrödinger, New York, NY) using the same force field and water model as for the monomer MD simulation. Dimer conformations were created by placing two copies of the representative monomer, cluster 1 in Fig. S1, at 4 nm center of mass (CoM) distance. Two configurations were created, parallel and orthogonal (90° rotation between the two monomers with respect to the long peptide axes). The dimers were then solvated, neutralized, and kept at 150 mM NaCl concentration after which they underwent 50 ns NPT simulation to relax the system. They were then submitted for 4 μs MD simulation on Anton.

We monitored secondary structure dynamics according to the method developed by Thirumalai’s group (75). Briefly, if the dihedral angles from two consecutive residues satisfy the definition of an α-helix (−80° ≤ ϕ ≤ −48° and −59°≤ ψ ≤ −27°) or β-strand (−150° ≤ ϕ ≤ −90° and 90 ≤ ψ ≤ 150°), the structures are considered to be α or β conformations, respectively. The changes of secondary structure over time are monitored by, 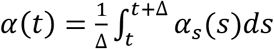 and 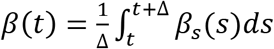, where t =s and Δ=1 ns. When the residues adopt α- or β-conformations, the *δ*_*i*,*α*_ = 1 or *δ*_*i*,*β*_ = 1.

### Accelerated MD simulations

To further extend conformational sampling, the resulting structures from the MD simulations on Anton were subjected to the accelerated MD (aMD) simulation method. The simulation procedure was adopted from the description by Pierce *et al.* (76) and the website (URL: http://ambermd.org/tutorials/advanced/tutorial22/) using Amber 14 software package (77). Briefly, dimer conformations from the last frame of the MD simulation on Anton and the from the two lowest energy minima were solvated, neutralized, and kept at 150 mM NaCl before being submitted for 500 ns aMD simulations. Simulations utilized the same force field and water model as previous simulations.

The principal component analysis of backbone dihedrals (dPC) (78) was used to calculate the energy landscape and identify the representative structures of the minima. The Fortran program (78) written by Dr. Yuguang Mu was used to perform this analysis. Intra-peptide contact probability maps were generated based on Cα atom contacts within the monomers using the GROMACS *mdmat* analysis tool.

### Monte Carlo pulling simulations

The Monte Carlo pulling (MCP) method was performed to simulate AFM force spectroscopy experiments using our previously described procedure (58) and a modified PROFASI package (79). Briefly, the two Cα of the N-terminal Cys residues of each monomer were defined as the pulling groups. A virtual spring was attached onto each pulling group and used to stretch them during the pulling process. The energy dynamics of the spring were calculated using the A2A spring function (PROFASI package) with the total energy during the course of pulling described by,

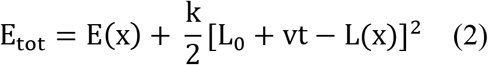

where *E(x)* describes the energy without an external force, *k* and *t* are the spring constant of the virtual spring, and *L*_0_ is the initial distance between the two Cα atoms. *L(x)* represents the real-time distance between the Cα atoms while *x* denotes the protein conformation being probed. When *v* = 0.083 fm per MC step, it mimics the pulling speed of 500 nm/s; which was used for all MCP simulations.

## Results

### Aβ40 Monomer Structure

We performed all-atom MD simulations of Aβ40 monomers to identify the most representative monomer structure. We adopted the approach from our recent simulations of the Aβ42 dimer (69). Briefly, the Aβ40 monomer structure was simulated for 500 ns, the most representative structure was then identified using cluster analysis by calculating the RMSD of backbone atoms between all pairs of structures with a cut-off at 3Å (80). The results of the cluster analysis are shown in Figure S1. Twelve clusters were identified, with the 1^st^ cluster comprising 47.5% of the entire population. The representative structure of this cluster contains a large α-helical segment in the central region of the peptide and is otherwise unstructured. Two copies of this structure were used to characterize the dimer conformation.

### Characterization of Aβ40 Dimer Formation

Two dimer systems were generated by placing copies of the monomer structure in orthogonal (90°) or parallel orientations, with respect to the long peptide axis, at 4 nm CoM distance, Figure 1 right column. Both dimer conformations were then simulated for 4 μs on the special purpose Anton supercomputer.

**Figure 1.**
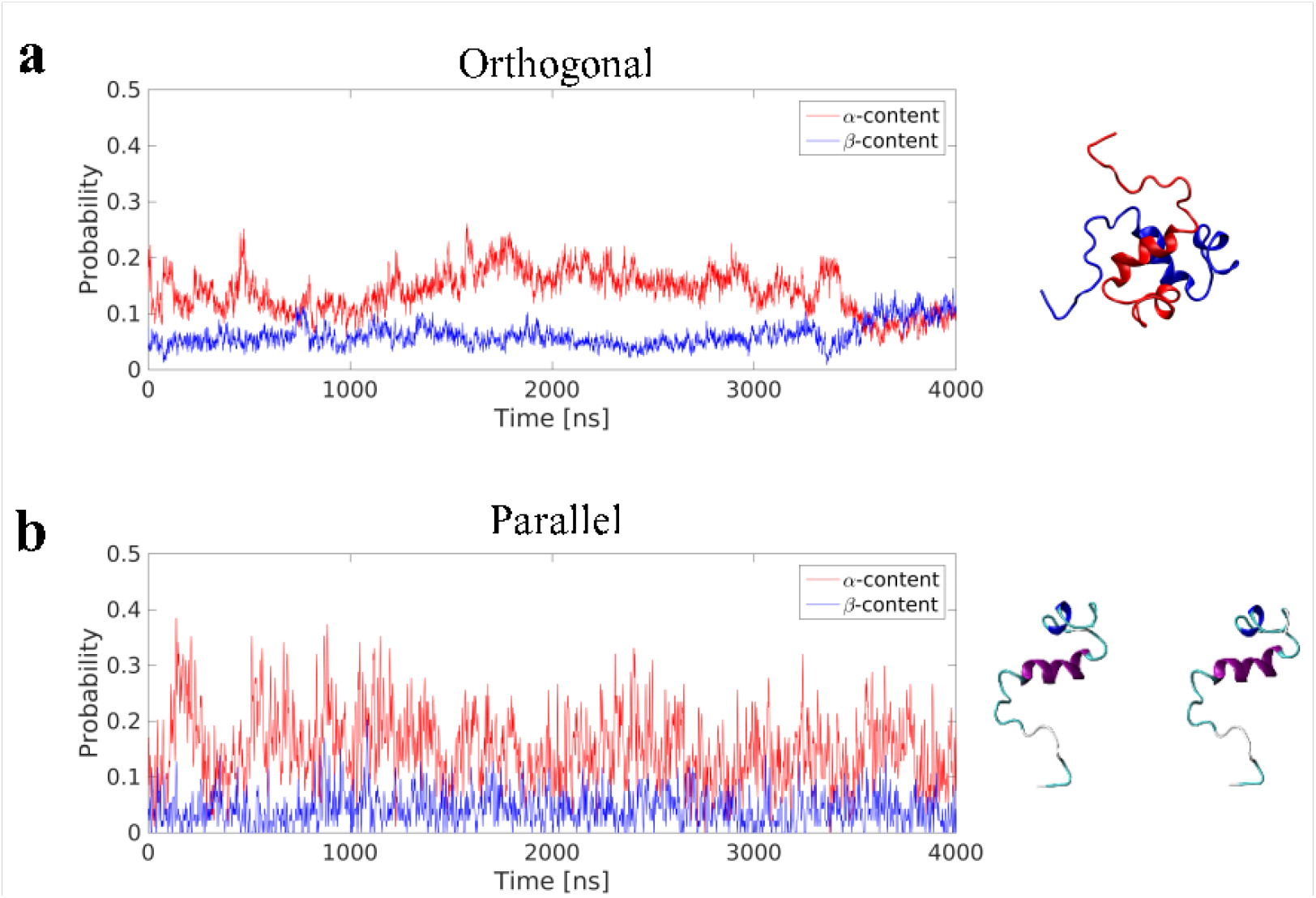
Time-resolved change in protein secondary structure during 4 μs all-atom MD simulations of Aβ40 dimers. Data for Aβ40 dimer in the orthogonal (a) and parallel (b) starting configuration are presented; right column shows a snapshot of the initial structures for each system.

To determine if the dimer simulations had converged, we monitored the time-dependent change in secondary structure of the peptides, Figure 1 left column. The graphs show that for the orthogonal configuration, α-helical content fluctuates with a decreasing tendency up to the 1 μs mark, after which the helical portion increases over the next 1 μs span, Figure 1a. Meanwhile, the β-content remains stable at approximately 5%, with minor fluctuations, until approximately 3.5 μs; after which a conversion from α-helical to β-structure is observed, with β-content reaching a maximum of ~12% at the end of the simulation. For the parallel configuration on the other hand, both α-helical to β-structure content fluctuate throughout the simulation, with averages of approximately 15% and 5%, respectively, Figure 1b. This suggests that, for both configurations, a local equilibrium state has not been reached.

We then used dPC analysis to analyze the energy landscape of the dimer. For both dimer configurations, several distinct energy minima were found, Figure S2Error! Reference source not found‥ Furthermore, both configurations show a rough and discontinuous energy landscape. This, in combination with the time-resolved change in secondary structure, suggests that the dimers are trapped in local energy minima, leading to insufficient sampling of the conformational space. To overcome this problem and to enhance the sampling of the conformational space, we extended the dimer simulation using aMD simulations (see specifics in Methods) allowing us to reach sampling enhancement by several orders of magnitude (76).

### Accelerated Molecular Dynamics Simulations of Dimers

The result of the aMD simulations for the dimer is presented in Figure S3. Several well-defined and separated energy minima were identified for the orthogonal system, Figure S3a, while the parallel system only has few energy minima that are clustered in the same region of the energy landscape, Figure S3b. The aMD results were then pooled and the concatenated data set underwent dPC analysis again, Figure 2a. The snapshots in the figure depict representative structures from the two lowest energy minima. It is evident, that the dimer does not adopt long β-structures but has a mixture of short helices and β-structures.

**Figure 2.**
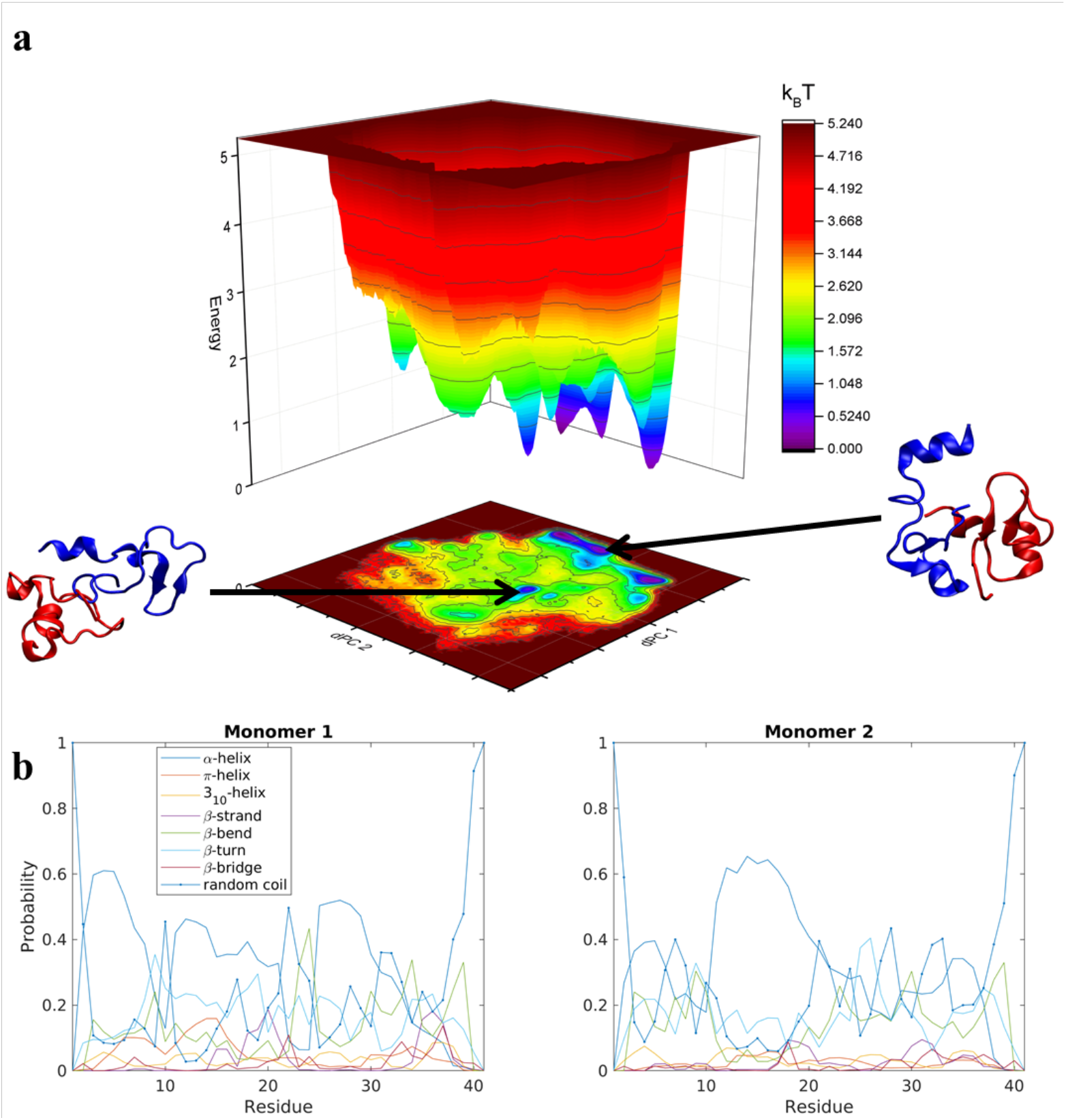
Analysis of Aβ40 dimers obtained from 3 μs aggregate accelerated MD simulations. (a) Free energy landscape based on dihedral principle component analysis of Aβ40 dimers; the two lowest energy structures are shown as cartoons. Blue depict Monomer 1 while red is Monomer 2. (b) Probability of each secondary structure type, determined by DSSP, for each monomer within the Aβ40 dimer, on a per residue basis.

The secondary structure of the dimers was characterized using DSSP (81). Each monomer was investigated separately with the results being displayed as residue specific probabilities, Figure 2b. Monomer 1 shows greater than 40% propensity for helix formation in residues 3-7, 11-13, and 25-29. β-structures are overall less likely compared to helices, however regions 10-30 and 35-38 have on average greater than 20% chance of β-structures. Monomer 2 on the other hand is more diverse, the helix probability is localized around residues 11-20, while collectively β-structures are more probable in the N- and C-terminal segments in residues 3-10 and 21-38, respectively.

To analyze the conformational diversity of the dimers we performed cluster analysis using the pooled aMD data. Similar to the analysis performed for monomers, clustering was performed using RMSD of backbone atoms between all pairs of structures with a cut-off at 3Å. Representative structures for the first 20 clusters are depicted in cartoon representation and relative populations on Figure 3. Structurally the clusters, with few exceptions, exhibit similar trends of low α-helical and β-structural content and high degree of unstructured regions. The main difference within the clusters arise from the different configurations of monomers.

**Figure 3.**
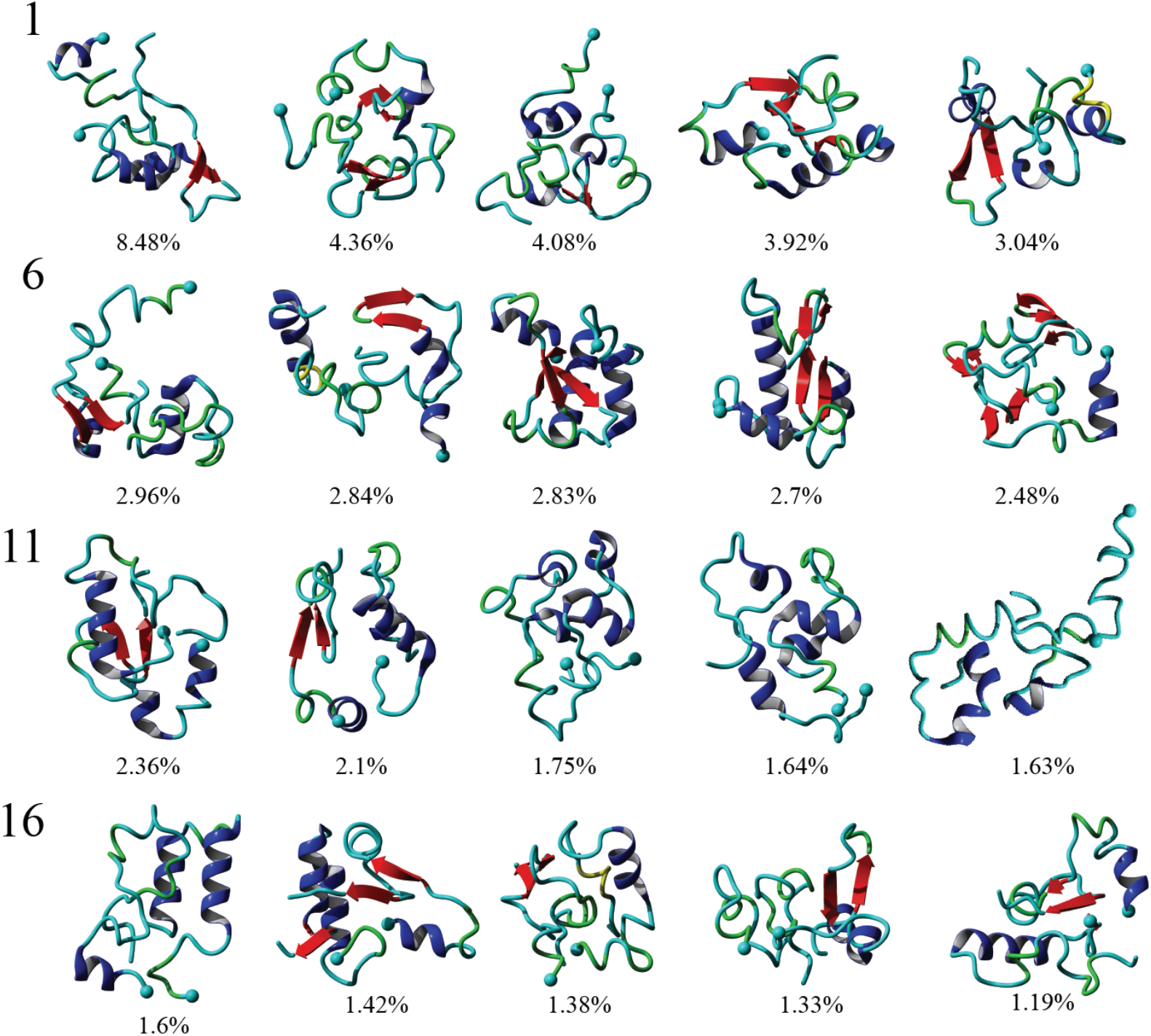
Cluster analysis of Aβ40 dimers obtained from 3 μs aggregate accelerated MD simulations. Representative structures of the top 20 clusters formed by Aβ40 dimers are presented with relative populations, as percent, for each cluster displayed below each structure. α-helices are colored blue while β-strands are in red. A solid sphere depicts the N-terminal Cα.

To identify segments important for the interaction of Aβ40 monomers, we performed analysis of the pair-wise residue interactions. Intra-peptide contact probability maps were generated based on Cα atom contacts within the monomers, Figure S4. For Monomer 1, interactions in three segments stand out, residues 5-12, residues 16-23, and residues 30-40, Figure S4a. The interactions within these three segments reveal that the monomer during the simulations, with high probability, is found in a compact turn-like conformation with C-terminal interacting with the central segment of the peptide. Monomer 2 on the other hand is more dynamic with few residues interacting within the N-terminal region and the 16-23 segment, Figure S4b. The interaction patterns of the two monomers reveal that, apart from neighbor residue interactions, the main difference is found in the way the two monomers interact with the 16-23 region; for Monomer 1 the interaction happens with residues 33-38, while for Monomer 2 it is residue 28-32, Figure 4a.

**Figure 4.**
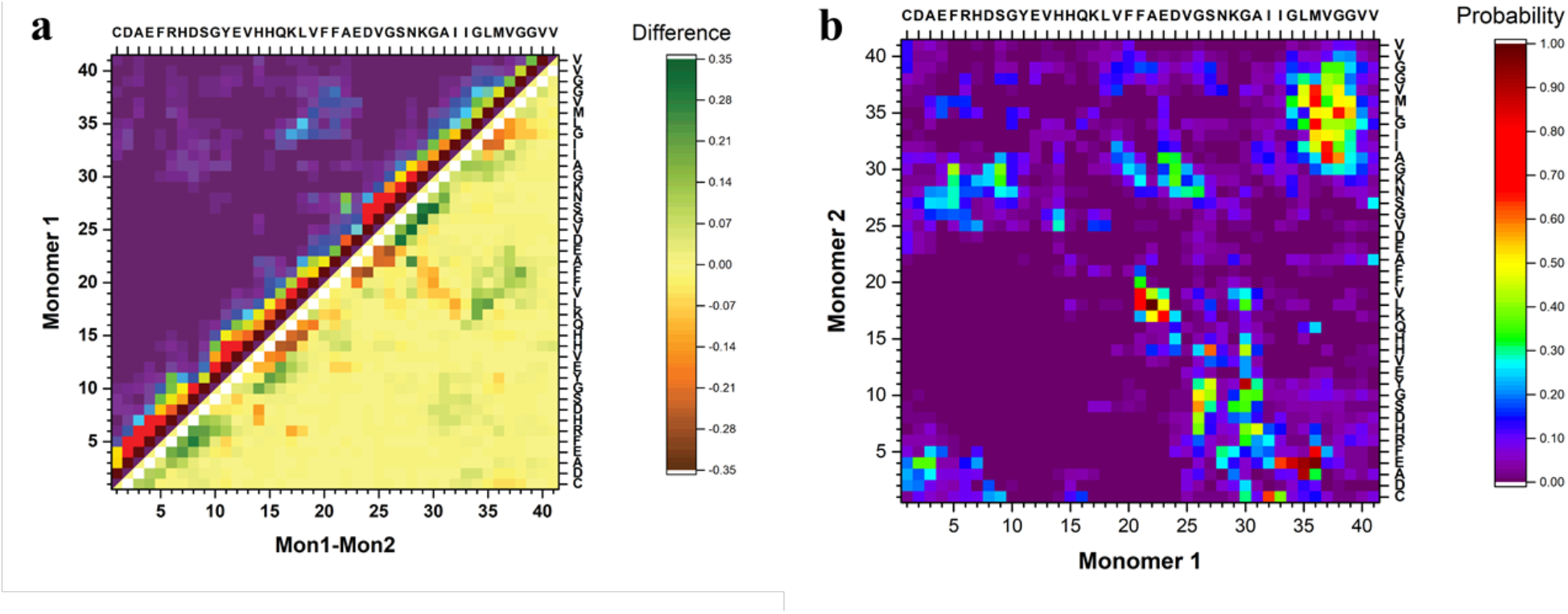
Analysis of inter-peptide interactions of Aβ40 dimers from 3 μs aggregate accelerated MD. (a) The difference in the contact probability between the two monomers and (b) the inter-peptide contact probability map for Cα atoms of dimers.

The inter-peptide interactions of the dimer were obtained using the pair-wise interactions of Cα atom between the monomers, Figure 4b. The contact map reveals that the interactions between the two monomers occur in the central region of the peptide as well as between the N-and C-terminals and the two C-termini. Comparison of the contact data and the dimer structures, revealed by cluster analysis, Figure 3, shows that the 20 most populated clusters are a mixture of different conformations that all contain N-C terminal interactions, with a few configurations also containing C-C terminal interactions. Monomer 1 primarily interacts through its central and C-terminal segments, while Monomer 2 interacts through the N- and C-terminal regions.

#### 1.1.1 Validation of Dimer Conformations

To validate the simulation results as well as identify the experimentally relevant conformations we used the Monte Carlo pulling approach to simulate AFM pulling experiments and to compare the simulated results with the experimental data. The rupture force and interaction patterns for the top candidates are presented in Figure 5. The interaction patterns of the simulated dissociation processes were normalized with respect to the experimentally obtained contour lengths. Experimentally observed values for the dissociation force was 56.6 ± 20.5 pN (STD), approximated using a Gaussian distribution, with a two-peak distribution of the interaction pattern favoring interaction in the N-terminal and central regions (47).

**Figure 5.**
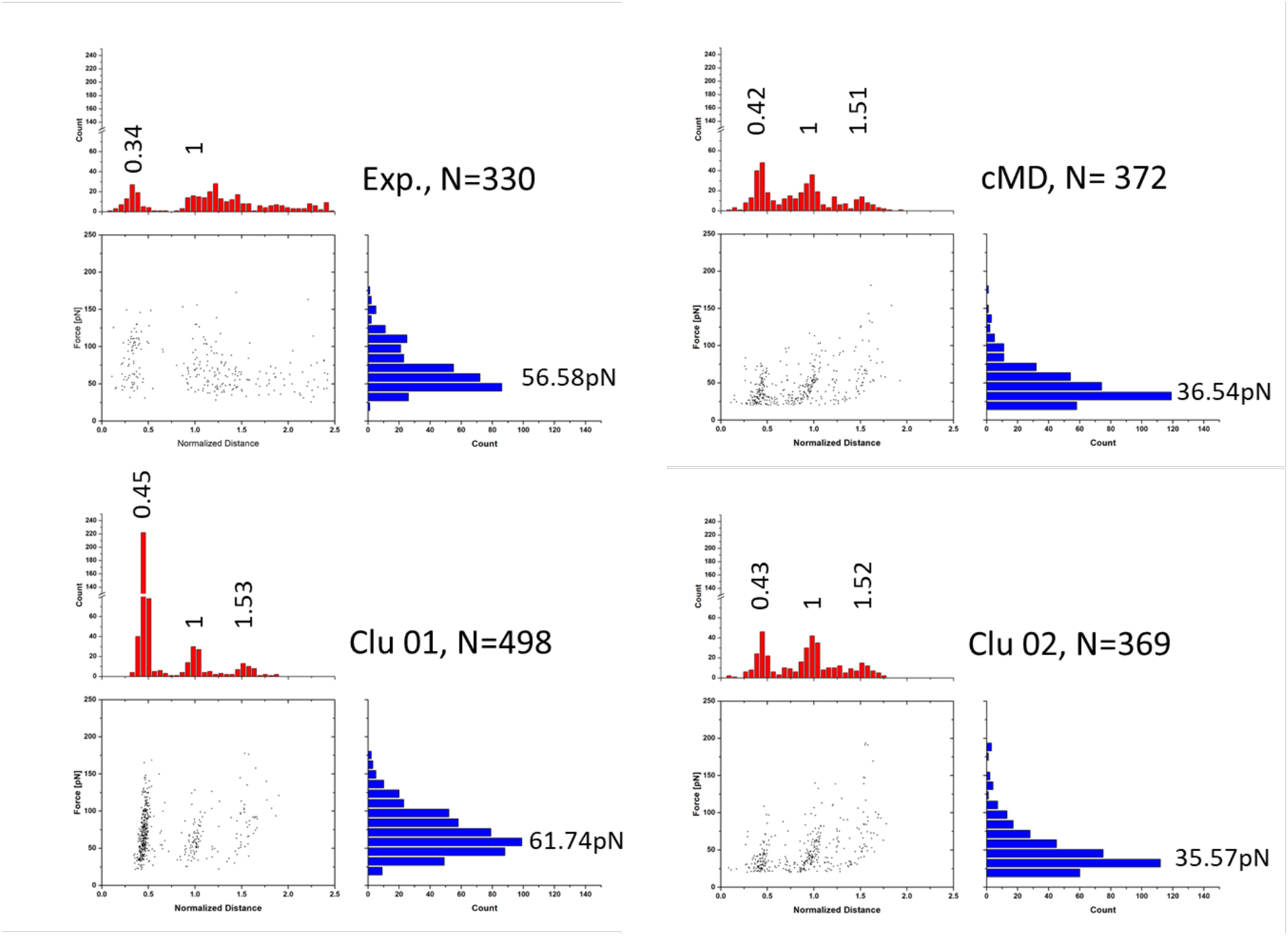
Force-induced dissociation results for Aβ40 dimers obtained from experiment (from (47)) and MCP simulations. Each dataset shows a scatter plot of Normalized Distance vs Force, a histogram of Force (blue), and a histogram of Normalized Distance; normalization was performed based on the experimentally observed contour lengths. Peak values, obtained using Gaussian distribution function, are presented above each peak of the histogram. Clu 01 and 02 are conformations from Figure 2, while “MD” is the most populated cluster following MD simulations. Statistical analysis was performed using two-sample Kolmogorov-Smirnov test with 0.05 significance level; only Clu 01 was statistically similar to the experimental data set, with p>0.066.

The dimer obtained following analysis of the MD simulations on Anton, named “MD” on Figure 4, shows a distinct three-peak interaction pattern, with majority of interactions located in the N-terminal and central regions of the proteins, while the dissociation force is 36.5 ± 18.4 pN. Dimer conformations from the two most populated clusters (Clu 01 and Clu 02) from Figure 2 (following the aMD simulations) produce rupture forces of 61.7 ± 27.5 pN and 35.6 ± 17.7 pN, respectively. Similar to the MD dimer, the two aMD conformations produce the distinct three-peak interaction pattern. However, Clu 01 shows a very large C-terminal peak. The dissociation of dimer Clu 01 is statistically similar to the experimentally observed results, using a non-parametric two-sample Kolmogorov-Smirnov with 0.05 significance.

To characterize the interaction pattern and the dissociation force of a dimer (within fibrils) with high β-structure content, we created two dimer conformations from NMR structures of Aβ40 fibrils with different morphologies (PDB IDs: 2LMN (wild-type) and 2MVX (Osaka mutant)). The dissociation patterns for the two fibril dimers are significantly different compared to experimental results and the results obtained for the MD and aMD dimers, Figure S6. Although, the fibril dimers contain the three-peak interaction pattern, the patterns are significantly different; for the 2LMN dimer the majority of interactions happen within the central part of the dimers, while for 2MVX dimer the interactions are dominated by the N- and C-terminals.

## 1.2 Discussion

Although the behavior of Aβ peptides have been subject to numerous studies, our present study presents a number of new features about the Aβ40 dimers. The equilibrated monomer structure, used as the initial conformation to characterize the dimerization process, is in line with recent data obtained using NMR and simulations of the Aβ proteins, which showed that the monomer has unstructured segments and can assume helical secondary structure (10, 82). Another interesting feature of the monomer structure is the presence of a turn on each side of the central helix, the turn conformation is believed to be the first folding event in the structural transition of Aβ proteins and important for the aggregation process (5, 83, 84).

Our computational analysis of the aggregation of Aβ40 into dimers reveal a broad range of peptide structures and very dynamic feature of the dimers. In particular, we did not identify significant β-conformation in the monomers within the dimer, Figure 3. The interaction of two monomers lead to conformational transitions within the monomers, accompanied by change in local structure of the peptides, leading to the formation of a stable dimer. Investigation of the dimer structures showed that the Aβ40 dimers exhibit a heterogeneous ensemble of conformations that contain a diverse number of structures. Dimers are primarily stabilized by interactions in the N-terminal region (residues 5-12), in the central hydrophobic region (residues 16-23), and in the C-terminal region (residues 30-40); with inter-peptide interactions focused around the N- and C- terminals. The 20 most populated clusters are a mixture of different conformations that all contain N-C terminal interactions, with a few configurations also containing C-C terminal interactions. Similar observations regarding the interaction pattern of Aβ40 dimers have been presented by Tarus *et al.* (85). The authors showed that regions, identified in our simulations, were also interacting and important for the stability of the dimer. However, unlike the dimer conformations identified here, their dimers contained significant β-structure content. More recent findings from the same group (86) show that the dimers structures are more diverse and do not contain a large extent of β-structure, and that the dimer is stabilized by nonspecific interactions. The low β-structure content is in agreement with our findings, and also can explain the role of structural plasticity in the interactions of Aβ oligomers with binding partners and ultimately their toxicity. The structural flexibility of the dimer may also play a role in the aggregation progression, where the free energy cost of transitioning from less ordered states is much less compared to dimeric states with high level of ordered β-structures.

We validated the dimer conformations using MCP approach to simulate the force-induced dissociation of the dimers and compared the obtained force and interaction patterns with experimental results. The simulations were performed at conditions identical to the experimental ones and allowed us to identify the dimer conformation of Clu 01 as the most probable dimer probed during experiments. Probing of dimer conformations with high degree of β-structure content, adopted from fibril structures, showed that such dimers produce dissociation forces significantly different compared to experiment as well as our simulated dimers. Furthermore, the interaction pattern of high β-content dimers was strongly shifted compared to experiments.

Comparing the Aβ40 dimer with the Aβ42 dimer, analyzed in our recent publication (69), shows that the Aβ42 dimer is stabilized by interactions in the central region (residues 16-23) between the two monomers as well as C-C terminal interactions through residues 30-36 and 36-42. Interactions also occur between the N-termini of the two monomers. Suggesting that the two extra C-terminal amino acids of Aβ42 affects the spatial orientation within the dimer as well as the inter-peptide interaction pattern of the monomers. These finding are in line with recent finding about the monomeric Aβ peptides (82), which show that while the two alloforms show similar structural elements, their conformations are different and that in turn has a large effect on the inter-molecular interactions of the peptides.

## 1.3 Conclusions

All-atom MD simulations allowed us to structurally characterize Aβ40 dimers. Structures were organized in clusters, with ~54% represented in the 20 most populated clusters. Dimers are stabilized by interactions in the central hydrophobic region (residues 17-21) as well as N-C terminal interactions (residues 1-10 and 30-40), through hydrophobic interactions and H-bonds. Aβ40 dimer did not show parallel in-register β-sheet structures, as one may expect based on the known structures of Aβ fibrils. Comparison of Aβ40 to Aβ42 dimers revealed differences in their conformations. Aβ40 dimers are stabilized primarily by interactions within the central hydrophobic regions and the N-terminal regions, whereas Aβ42 dimers are stabilized by interactions in the central and C-terminal regions. Aβ40 dimers are more dynamic compared to Aβ42 dimers. Comparison, based on MCP simulations, between Aβ40 and Aβ42 showed that overall, the dimers of both alloforms exhibit similar interaction strengths. However, the interaction maps, and more importantly the patterns, clearly show differences.

## List of abbreviations

AD: Alzheimer’s disease
AFM: Atomic force microscopy
aMD: Accelerated MD
CoM: Center of mass
dPC: Dihedral principal component
DSSP: Define secondary structure of proteins
FRET: Förster resonance energy transfer
MCP: Monte Carlo pulling
MD: Molecular dynamics
NPT: Isothermal-isobaric
PDB: Protein data bank
RMSD: root-mean square deviation
TAPIN: Tethered approach for probing inter-molecular interactions

## Acknowledgments

This work was supported by NIH grants GM096039 and GM100156, NSF grant 1004094, and the PSCA14025P award for computer time on Anton at Pittsburgh Supercomputing Center (PSC) - all to YLL. MH was supported by the Bukey Memorial Fellowship. Computational modeling was performed using facilities of the Holland Computing Center at the University of Nebraska (supported by the Nebraska Research Initiative) and the San Diego Supercomputing Center at the University of California San Diego through the Extreme Science and Engineering Discovery Environment (XSEDE; supported by National Science Foundation [ACI-1053575 for XSEDE]). Anton computer time was provided by the PSC through Grant R01GM116961 from the NIH. The Anton machine at PSC was generously made available by D.E. Shaw Research. We thank Dr. Yuguang Mu for providing the dPCA Fortran program.

## Author contributions

MH, YZ, and YLL conceived and designed the overall study. MH and YZ performed the Aβ40 simulations and analyzed all the simulation data. Z.L. performed the analysis of the experimental dataset. All authors discussed the results and wrote the paper.

## Additional information

Supplementary Information accompanies this paper at: http://

## References

1. Dobson, C. M. (2003) Protein folding and misfolding. Nature 426, 884–890

2. Chiti, F., and Dobson, C. M. (2006) Protein Misfolding, Functional Amyloid, and Human Disease. Annu. Rev. Biochem. 75, 333–366

3. Petkova, A. T., Leapman, R. D., Guo, Z., Yau, W.-M., Mattson, M. P., and Tycko, R. (2005) Self-Propagating, Molecular-Level Polymorphism in Alzheimer’s ß-Amyloid Fibrils. Science 307, 262–265

4. Hardy, J., and Selkoe, D. J. (2002) The amyloid hypothesis of Alzheimer’s disease: progress and problems on the road to therapeutics. Science 297, 353–356

5. Lührs, T., Ritter, C., Adrian, M., Riek-Loher, D., Bohrmann, B., Döbeli, H., Schubert, D., and Riek, R. (2005) 3D structure of Alzheimer’s amyloid-β(1–42) fibrils. Proc. Natl. Acad. Sci. USA 102, 17342–17347

6. Xiao, Y., Ma, B., McElheny, D., Parthasarathy, S., Long, F., Hoshi, M., Nussinov, R., and Ishii, Y. (2015) Aβ(1-42) fibril structure illuminates self-recognition and replication of amyloid in Alzheimer’s disease. Nat. Struct. Mol. Biol. 22, 499–505

7. Walti, M. A., Ravotti, F., Arai, H., Glabe, C. G., Wall, J. S., Bockmann, A., Guntert, P., Meier, B. H., and Riek, R. (2016) Atomic-resolution structure of a disease-relevant Abeta(1-42) amyloid fibril. Proc Natl Acad Sci U S A 113, E4976–4984

8. Crescenzi, O., Tomaselli, S., Guerrini, R., Salvadori, S., D’Ursi, A. M., Temussi, P. A., and Picone, D. (2002) Solution structure of the Alzheimer amyloid β-peptide (1–42) in an apolar microenvironment. Euro. J. Biochem. 269, 5642–5648

9. Sgourakis, N. G., Merced-Serrano, M., Boutsidis, C., Drineas, P., Du, Z., Wang, C., and Garcia, A. E. (2011) Atomic-Level Characterization of the Ensemble of the Aβ(1–42) Monomer in Water Using Unbiased Molecular Dynamics Simulations and Spectral Algorithms. J. Mol. Biol. 405, 570–583

10. Vivekanandan, S., Brender, J. R., Lee, S. Y., and Ramamoorthy, A. (2011) A partially folded structure of amyloid-beta(1-40) in an aqueous environment. Biochem. Biophys. Res. Commun. 411, 312–316

11. Glabe, C. G. (2008) Structural classification of toxic amyloid oligomers. The Journal of biological chemistry 283, 29639–29643

12. Glabe, C. G. (2006) Common mechanisms of amyloid oligomer pathogenesis in degenerative disease. Neurobiol. Aging 27, 570–575

13. Yu, L., Edalji, R., Harlan, J. E., Holzman, T. F., Lopez, A. P., Labkovsky, B., Hillen, H., Barghorn, S., Ebert, U., Richardson, P. L., Miesbauer, L., Solomon, L., Bartley, D., Walter, K., Johnson, R. W., Hajduk, P. J., and Olejniczak, E. T. (2009) Structural Characterization of a Soluble Amyloid β-Peptide Oligomer. Biochemistry 48, 1870–1877

14. Laganowsky, A., Liu, C., Sawaya, M. R., Whitelegge, J. P., Park, J., Zhao, M., Pensalfini, A., Soriaga, A. B., Landau, M., Teng, P. K., Cascio, D., Glabe, C., and Eisenberg, D. (2012) Atomic view of a toxic amyloid small oligomer. Science 335, 1228–1231

15. Liu, P., Reed, M. N., Kotilinek, L. A., Grant, M. K., Forster, C. L., Qiang, W., Shapiro, S. L., Reichl, J. H., Chiang, A. C., Jankowsky, J. L., Wilmot, C. M., Cleary, J. P., Zahs, K. R., and Ashe, K. H. (2015) Quaternary Structure Defines a Large Class of Amyloid-beta Oligomers Neutralized by Sequestration. Cell Rep. 11, 1760–1771

16. Zhang, Y., McLaughlin, R., Goodyer, C., and LeBlanc, A. (2002) Selective cytotoxicity of intracellular amyloid beta peptide 1-42 through p53 and Bax in cultured primary human neurons. J Cell Biol 156, 519–529

17. Rajasekhar, K., Chakrabarti, M., and Govindaraju, T. (2015) Function and toxicity of amyloid beta and recent therapeutic interventions targeting amyloid beta in Alzheimer’s disease. Chem Commun (Camb) 51, 13434–13450

18. Nagy, Z., Esiri, M. M., and Smith, A. D. (1998) The cell division cycle and the pathophysiology of Alzheimer’s disease. Neuroscience 87, 731–739

19. Snyder, E. M., Nong, Y., Almeida, C. G., Paul, S., Moran, T., Choi, E. Y., Nairn, A. C., Salter, M. W., Lombroso, P. J., Gouras, G. K., and Greengard, P. (2005) Regulation of NMDA receptor trafficking by amyloid-beta. Nat Neurosci 8, 1051–1058

20. Shankar, G. M., Bloodgood, B. L., Townsend, M., Walsh, D. M., Selkoe, D. J., and Sabatini, B. L. (2007) Natural oligomers of the Alzheimer amyloid-beta protein induce reversible synapse loss by modulating an NMDA-type glutamate receptor-dependent signaling pathway. J Neurosci 27, 2866–2875

21. Hsieh, H., Boehm, J., Sato, C., Iwatsubo, T., Tomita, T., Sisodia, S., and Malinow, R. (2006) AMPAR removal underlies Abeta-induced synaptic depression and dendritic spine loss. Neuron 52, 831–843

22. Serra-Batiste, M., Ninot-Pedrosa, M., Bayoumi, M., Gairi, M., Maglia, G., and Carulla, N. (2016) Abeta42 assembles into specific beta-barrel pore-forming oligomers in membrane-mimicking environments. Proc Natl Acad Sci U S A 113, 10866–10871

23. Lashuel, H. A., Hartley, D., Petre, B. M., Walz, T., and Lansbury, P. T., Jr. (2002) Neurodegenerative disease: amyloid pores from pathogenic mutations. Nature 418, 291

24. Tougu, V., Tiiman, A., and Palumaa, P. (2011) Interactions of Zn(II) and Cu(II) ions with Alzheimer’s amyloid-beta peptide. Metal ion binding, contribution to fibrillization and toxicity. Metallomics 3, 250–261

25. Ramamoorthy, A., and Lim, M. H. (2013) Structural characterization and inhibition of toxic amyloid-beta oligomeric intermediates. Biophys J 105, 287–288

26. LaFerla, F. M., Green, K. N., and Oddo, S. (2007) Intracellular amyloid-beta in Alzheimer’s disease. Nat Rev Neurosci 8, 499–509

27. Choi, J. S., Braymer, J. J., Nanga, R. P., Ramamoorthy, A., and Lim, M. H. (2010) Design of small molecules that target metal-A{beta} species and regulate metal-induced A{beta} aggregation and neurotoxicity. Proc Natl Acad Sci U S A 107, 21990–21995

28. Kayed, R., Pensalfini, A., Margol, L., Sokolov, Y., Sarsoza, F., Head, E., Hall, J., and Glabe, C. (2009) Annular protofibrils are a structurally and functionally distinct type of amyloid oligomer. The Journal of biological chemistry 284, 4230–4237

29. Harper, J. D., Lieber, C. M., and Lansbury, P. T. (1997) Atomic force microscopic imaging of seeded fibril formation and fibril branching by the Alzheimer’s disease amyloid-β protein. Chemistry & Biology 4, 951–959

30. Ahmed, M., Davis, J., Aucoin, D., Sato, T., Ahuja, S., Aimoto, S., Elliott, J. I., Van Nostrand, W. E., and Smith, S. O. (2010) Structural conversion of neurotoxic amyloid-beta(1-42) oligomers to fibrils. Nat Struct Mol Biol 17, 561–567

31. Ono, K., Condron, M. M., and Teplow, D. B. (2009) Structure-neurotoxicity relationships of amyloid beta-protein oligomers. Proc Natl Acad Sci U S A 106, 14745–14750

32. Sarkar, B., Mithu, V. S., Chandra, B., Mandal, A., Chandrakesan, M., Bhowmik, D., Madhu, P. K., and Maiti, S. (2014) Significant structural differences between transient amyloid-beta oligomers and less-toxic fibrils in regions known to harbor familial Alzheimer’s mutations. Angew Chem Int Ed Engl 53, 6888–6892

33. Bhowmik, D., Mote, K. R., MacLaughlin, C. M., Biswas, N., Chandra, B., Basu, J. K., Walker, G. C., Madhu, P. K., and Maiti, S. (2015) Cell-Membrane-Mimicking Lipid-Coated Nanoparticles Confer Raman Enhancement to Membrane Proteins and Reveal Membrane-Attached Amyloid-β Conformation. ACS Nano 9, 9070–9077

34. Tomita, T., Maruyama, K., Saido, T. C., Kume, H., Shinozaki, K., Tokuhiro, S., Capell, A., Walter, J., Grunberg, J., Haass, C., Iwatsubo, T., and Obata, K. (1997) The presenilin mutation (N141I) linked to familial Alzheimer disease (Volga German families) increases the secretion of amyloid beta protein ending at the 42nd (or 43rd) residue. Proc Natl Acad Sci U S A 94, 2025–2030

35. O’Nuallain, B., Shivaprasad, S., Kheterpal, I., and Wetzel, R. (2005) Thermodynamics of A beta(1-40) amyloid fibril elongation. Biochemistry 44, 12709–12718

36. Hellstrand, E., Boland, B., Walsh, D. M., and Linse, S. (2010) Amyloid beta-protein aggregation produces highly reproducible kinetic data and occurs by a two-phase process. ACS Chem Neurosci 1, 13–18

37. Vandersteen, A., Hubin, E., Sarroukh, R., De Baets, G., Schymkowitz, J., Rousseau, F., Subramaniam, V., Raussens, V., Wenschuh, H., Wildemann, D., and Broersen, K. (2012) A comparative analysis of the aggregation behavior of amyloid-beta peptide variants. FEBS Lett 586, 4088–4093

38. Murphy, M. P., and LeVine, H., 3rd. (2010) Alzheimer’s disease and the amyloid-beta peptide. J Alzheimers Dis 19, 311–323

39. Selkoe, D. J. (2001) Alzheimer’s disease: genes, proteins, and therapy. Physiol Rev 81, 741–766

40. Bibl, M., Gallus, M., Welge, V., Lehmann, S., Sparbier, K., Esselmann, H., and Wiltfang, J. (2012) Characterization of cerebrospinal fluid aminoterminally truncated and oxidized amyloid-beta peptides. Proteomics Clin Appl 6, 163–169

41. Bitan, G., Kirkitadze, M. D., Lomakin, A., Vollers, S. S., Benedek, G. B., and Teplow, D. B. (2003) Amyloid beta -protein (Abeta) assembly: Abeta 40 and Abeta 42 oligomerize through distinct pathways. Proc Natl Acad Sci U S A 100, 330–335

42. Orte, A., Birkett, N. R., Clarke, R. W., Devlin, G. L., Dobson, C. M., and Klenerman, D. (2008) Direct characterization of amyloidogenic oligomers by single-molecule fluorescence. Proceedings of the National Academy of Sciences 105, 14424–14429

43. Yu, H., Dee, D. R., Liu, X., Brigley, A. M., Sosova, I., and Woodside, M. T. (2015) Protein misfolding occurs by slow diffusion across multiple barriers in a rough energy landscape. Proceedings of the National Academy of Sciences 112, 8308–8313

44. Calamai, M., and Pavone, F. S. (2011) Single Molecule Tracking Analysis Reveals That the Surface Mobility of Amyloid Oligomers Is Driven by Their Conformational Structure. Journal of the American Chemical Society 133, 12001–12008

45. Brucale, M., Schuler, B., and Samorì, B. (2014) Single-Molecule Studies of Intrinsically Disordered Proteins. Chemical Reviews 114, 3281–3317

46. Kim, B.-H., Palermo, N. Y., Lovas, S., Zaikova, T., Keana, J. F. W., and Lyubchenko, Y. L. (2011) Single-Molecule Atomic Force Microscopy Force Spectroscopy Study of Aβ-40 Interactions. Biochemistry 50, 5154–5162

47. Lv, Z., Roychaudhuri, R., Condron, M. M., Teplow, D. B., and Lyubchenko, Y. L. (2013) Mechanism of amyloid beta-protein dimerization determined using single-molecule AFM force spectroscopy. Sci Rep 3, 2880

48. Krasnoslobodtsev, A. V., Volkov, I. L., Asiago, J. M., Hindupur, J., Rochet, J. C., and Lyubchenko, Y. L. (2013) alpha-Synuclein misfolding assessed with single molecule AFM force spectroscopy: effect of pathogenic mutations. Biochemistry 52, 7377–7386

49. Kim, B. H., and Lyubchenko, Y. L. (2014) Nanoprobing of misfolding and interactions of amyloid beta 42 protein. Nanomedicine 10, 871–878

50. Lovas, S., Zhang, Y., Yu, J., and Lyubchenko, Y. L. (2013) Molecular Mechanism of Misfolding and Aggregation of Aβ(13–23). J. Phys. Chem. B 117, 6175–6186

51. Banerjee, S., Sun, Z., Hayden, E. Y., Teplow, D. B., and Lyubchenko, Y. L. (2017) Nanoscale Dynamics of Amyloid β-42 Oligomers As Revealed by High-Speed Atomic Force Microscopy. ACS Nano 11, 12202–12209

52. Lv, Z., Krasnoslobodtsev, A. V., Zhang, Y., Ysselstein, D., Rochet, J.-C., Blanchard, S. C., and Lyubchenko, Y. L. (2015) Direct Detection of α-Synuclein Dimerization Dynamics: Single-molecule Fluorescence Analysis. Biophys J 108, 2038–2047

53. Lv, Z., Krasnoslobodtsev, A. V., Zhang, Y., Ysselstein, D., Rochet, J. C., Blanchard, S. C., and Lyubchenko, Y. L. (2016) Effect of acidic pH on the stability of alpha-synuclein dimers. Biopolymers 105, 715–724

54. Breydo, L., and Uversky, V. N. (2015) Structural, morphological, and functional diversity of amyloid oligomers. FEBS Lett 589, 2640–2648

55. Bemporad, F., and Chiti, F. (2012) Protein misfolded oligomers: experimental approaches, mechanism of formation, and structure-toxicity relationships. Chem Biol 19, 315–327

56. Baumketner, A., Bernstein, S. L., Wyttenbach, T., Bitan, G., Teplow, D. B., Bowers, M. T., and Shea, J. E. (2006) Amyloid beta-protein monomer structure: a computational and experimental study. Protein Sci 15, 420–428

57. Sgourakis, N. G., Yan, Y., McCallum, S. A., Wang, C., and Garcia, A. E. (2007) The Alzheimer’s peptides Abeta40 and 42 adopt distinct conformations in water: a combined MD / NMR study. J Mol Biol 368, 1448–1457

58. Zhang, Y., and Lyubchenko, Y. L. (2014) The structure of misfolded amyloidogenic dimers: computational analysis of force spectroscopy data. Biophys J 107, 2903–2910

59. Jang, S., and Shin, S. (2008) Computational study on the structural diversity of amyloid Beta Peptide (abeta(10-35)) oligomers. J Phys Chem B 112, 3479–3484

60. Fisher, C. K., Ullman, O., and Stultz, C. M. (2013) Comparative studies of disordered proteins with similar sequences: application to Abeta40 and Abeta42. Biophys J 104, 1546–1555

61. Flock, D., Colacino, S., Colombo, G., and Di Nola, A. (2006) Misfolding of the amyloid beta-protein: a molecular dynamics study. Proteins 62, 183–192

62. Yang, M., and Teplow, D. B. (2008) Amyloid beta-protein monomer folding: free-energy surfaces reveal alloform-specific differences. Journal of molecular biology 384, 450–464

63. Urbanc, B., Cruz, L., Yun, S., Buldyrev, S. V., Bitan, G., Teplow, D. B., and Stanley, H. E. (2004) In silico study of amyloid beta-protein folding and oligomerization. Proc Natl Acad Sci U S A 101, 17345–17350

64. Urbanc, B., Betnel, M., Cruz, L., Bitan, G., and Teplow, D. B. (2010) Elucidation of amyloid beta-protein oligomerization mechanisms: discrete molecular dynamics study. J Am Chem Soc 132, 4266–4280

65. Bernstein, S. L., Dupuis, N. F., Lazo, N. D., Wyttenbach, T., Condron, M. M., Bitan, G., Teplow, D. B., Shea, J. E., Ruotolo, B. T., Robinson, C. V., and Bowers, M. T. (2009) Amyloid-beta protein oligomerization and the importance of tetramers and dodecamers in the aetiology of Alzheimer’s disease. Nat Chem 1, 326–331

66. Zheng, W., Tsai, M.-Y., Chen, M., and Wolynes, P. G. (2016) Exploring the aggregation free energy landscape of the amyloid-β protein (1–40). Proceedings of the National Academy of Sciences 113, 11835–11840

67. Tarus, B., Tran, T. T., Nasica-Labouze, J., Sterpone, F., Nguyen, P. H., and Derreumaux, P. (2015) Structures of the Alzheimer’s Wild-Type Aβ1-40 Dimer from Atomistic Simulations. The Journal of Physical Chemistry B 119, 10478–10487

68. Watts, C. R., Gregory, A. J., Frisbie, C. P., and Lovas, S. (2017) Structural properties of amyloid β(1-40) dimer explored by replica exchange molecular dynamics simulations. Proteins: Structure, Function, and Bioinformatics 85, 1024–1045

69. Zhang, Y., Hashemi, M., Lv, Z., and Lyubchenko, Y. L. (2016) Self-assembly of the full-length amyloid Abeta42 protein in dimers. Nanoscale 8, 18928–18937

70. Shaw, D. E., Dror, R. O., Salmon, J. K., Grossman, J. P., Mackenzie, K. M., Bank, J. A., Young, C., Deneroff, M. M., Batson, B., Bowers, K. J., Chow, E., Eastwood, M. P., Ierardi, D. J., Klepeis, J. L., Kuskin, J. S., Larson, R. H., Lindorff-Larsen, K., Maragakis, P., Moraes, M. A., Piana, S., Shan, Y., and Towles, B. (2009) Millisecond-scale molecular dynamics simulations on Anton. In Proceedings of the Conference on High Performance Computing Networking, Storage and Analysis pp. 1–11, ACM, Portland, Oregon

71. Shaw, D. E., Maragakis, P., Lindorff-Larsen, K., Piana, S., Dror, R. O., Eastwood, M. P., Bank, J. A., Jumper, J. M., Salmon, J. K., Shan, Y., and Wriggers, W. (2010) Atomic-Level Characterization of the Structural Dynamics of Proteins. Science 330, 341–346

72. Hess, B., Kutzner, C., van der Spoel, D., and Lindahl, E. (2008) GROMACS 4: Algorithms for Highly Efficient, Load-Balanced, and Scalable Molecular Simulation. J. Chem. Theory Comput. 4, 435–447

73. Lindorff-Larsen, K., Piana, S., Palmo, K., Maragakis, P., Klepeis, J. L., Dror, R. O., and Shaw, D. E. (2010) Improved side-chain torsion potentials for the Amber ff99SB protein force field. Proteins 78, 1950–1958

74. Jorgensen, W. L., Chandrasekhar, J., Madura, J. D., Impey, R. W., and Klein, M. L. (1983) Comparison of simple potential functions for simulating liquid water. J. Chem. Phys. 79, 926–935

75. Reddy, G., Straub, J. E., and Thirumalai, D. (2009) Dynamics of locking of peptides onto growing amyloid fibrils. Proc. Natl. Acad. Sci. USA 106, 11948–11953

76. Pierce, L. C., Salomon-Ferrer, R., Augusto, F. d. O. C., McCammon, J. A., and Walker, R. C. (2012) Routine Access to Millisecond Time Scale Events with Accelerated Molecular Dynamics. J. Chem. Theory Comput. 8, 2997–3002

77. Case, D. A., Babin, V., Berryman, J. T., Betz, R. M., Cai, Q., Cerutti, D. S., Cheatham, T. E., Darden, T. A., Duke, R. E., Gohlke, H., Goetz, A. W., Gusarov, S., Homeyer, N., Janowski, P., Kaus, J., Kolossváry, I., Kovalenko, A., Lee, T. S., LeGrand, S., Luchko, T., Luo, R., Madej, B., Merz, K. M., Paesani, F., Roe, D. R., Roitberg, A., Sagui, C., Salomon-Ferrer, R., Seabra, G., Simmerling, C. L., Smith, W., Swails, J., Walker, Wang, J., Wolf, R. M., Wu, X., and Kollman, P. A. (2014) {Amber 14}

78. Mu, Y., Nguyen, P. H., and Stock, G. (2005) Energy landscape of a small peptide revealed by dihedral angle principal component analysis. Proteins 58, 45–52

79. Irback, A., and Mohanty, S. (2006) PROFASI: A Monte Carlo simulation package for protein folding and aggregation. J Comput Chem 27, 1548–1555

80. Daura, X., Gademann, K., Jaun, B., Seebach, D., van Gunsteren, W. F., and Mark, A. E. (1999) Peptide Folding: When Simulation Meets Experiment. Angew. Chem. Int. Ed. Engl. 38, 236–240

81. Touw, W. G., Baakman, C., Black, J., te Beek, T. A. H., Krieger, E., Joosten, R. P., and Vriend, G. (2015) A series of PDB-related databanks for everyday needs. Nucleic Acids Res 43, D364–D368

82. Roche, J., Shen, Y., Lee, J. H., Ying, J., and Bax, A. (2016) Monomeric Abeta(1-40) and Abeta(1-42) Peptides in Solution Adopt Very Similar Ramachandran Map Distributions That Closely Resemble Random Coil. Biochemistry 55, 762–775

83. Doran, T. M., Anderson, E. A., Latchney, S. E., Opanashuk, L. A., and Nilsson, B. L. (2012) An azobenzene photoswitch sheds light on turn nucleation in amyloid-beta self-assembly. ACS Chem Neurosci 3, 211–220

84. Lazo, N. D., Grant, M. A., Condron, M. C., Rigby, A. C., and Teplow, D. B. (2005) On the nucleation of amyloid beta-protein monomer folding. Protein Sci 14, 1581–1596

85. Tarus, B., Tran, T. T., Nasica-Labouze, J., Sterpone, F., Nguyen, P. H., and Derreumaux, P. (2015) Structures of the Alzheimer’s Wild-Type Abeta1-40 Dimer from Atomistic Simulations. J Phys Chem B 119, 10478–10487

86. Man, V. H., Nguyen, P. H., and Derreumaux, P. (2017) High-Resolution Structures of the Amyloid-beta 1-42 Dimers from the Comparison of Four Atomistic Force Fields. J Phys Chem B

