## Supplementary Information for "Spontaneous Self-assembly of Amyloid β (1-40) into Dimers"

**Running title**: Amyloid β dimer formation

**Keywords:** Amyloid Aβ40, Amyloid Aggregation, Dimer assembly, AFM force spectroscopy, MD simulations.

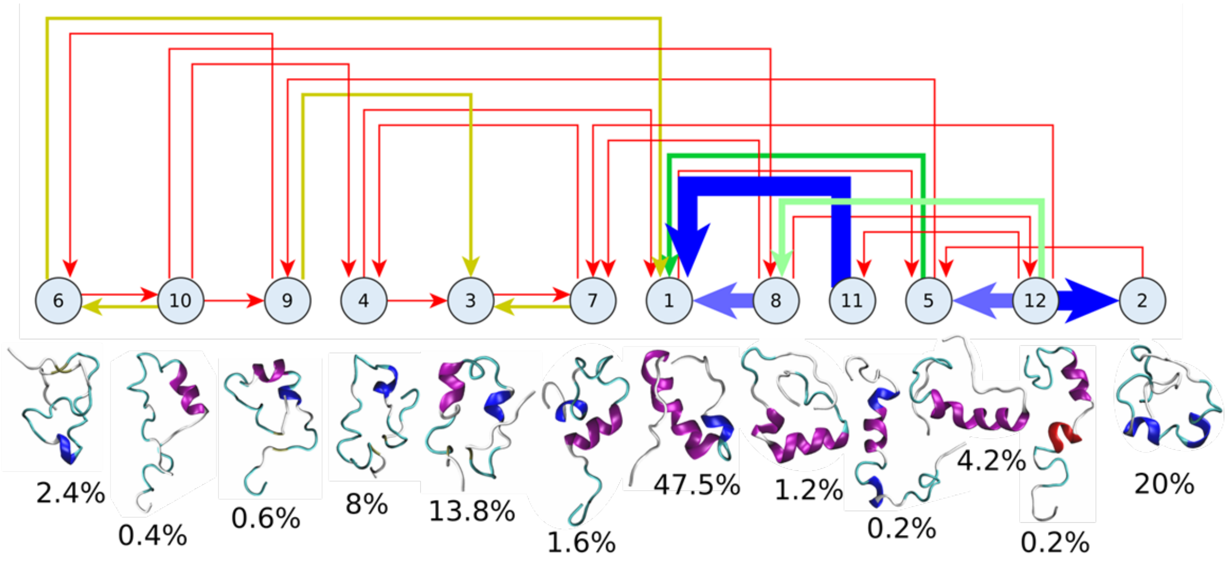


**Figure S1.** Cluster analysis of 500 ns conventional molecular dynamics simulation of Aβ40 monomer from PDB ID: 1AML. Representative structure for each cluster is shown below the cluster node together with the relative population percentage. Thickness of connecting links indicate the relative transition frequency. Proteins are shown in cartoon representation using the VMD secondary structure color scheme, with α-, π-, and 3/10-helices in purple, blue, and red respectively.

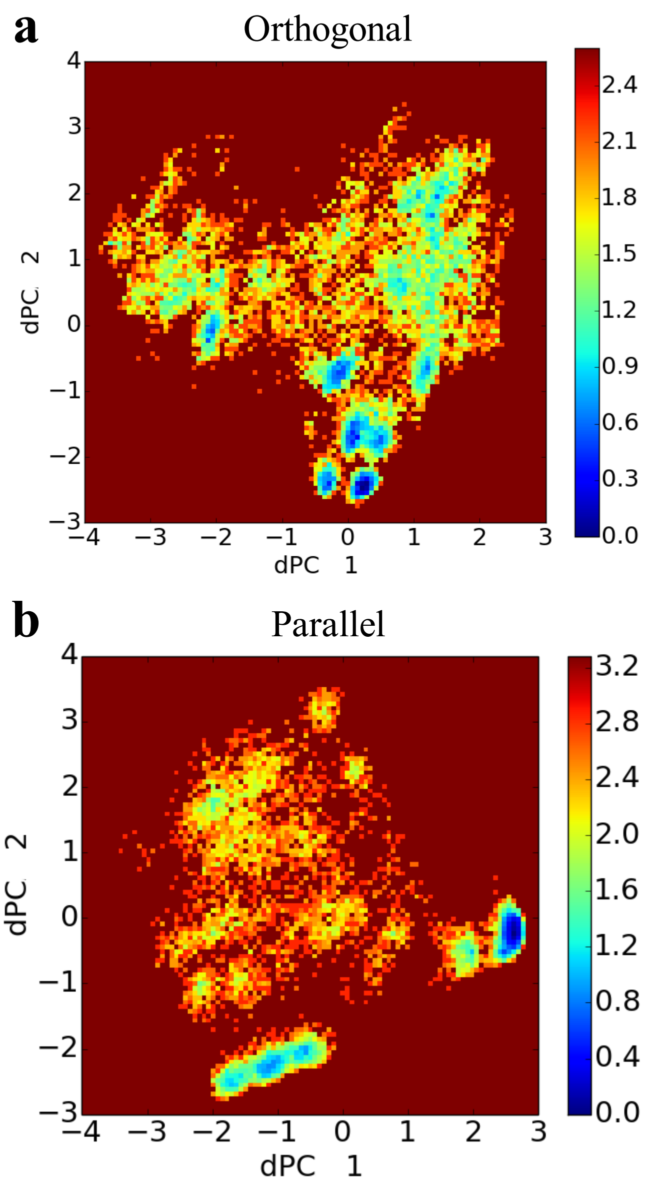


**Figure S2.** Free energy landscape of 4 μs MD simulations of Aβ40. Energy landscapes of Aβ40 in orthogonal (a) and parallel (b) configurations. Units of color bars are in $k_BT$.

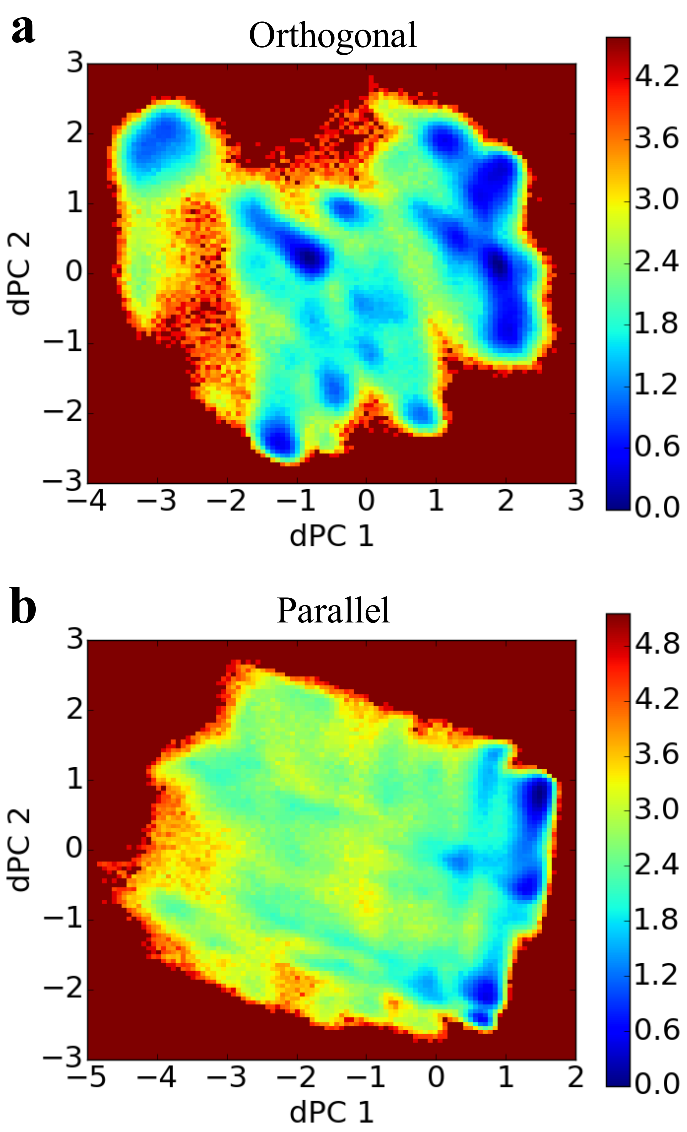


**Figure S3.** Free energy landscape of Aβ40 dimers from 1.5μs aggregate accelerated MD simulations. Aβ40 dimer in orthogonal (a) and parallel (b) configurations. Color bar units are in $k_BT$.

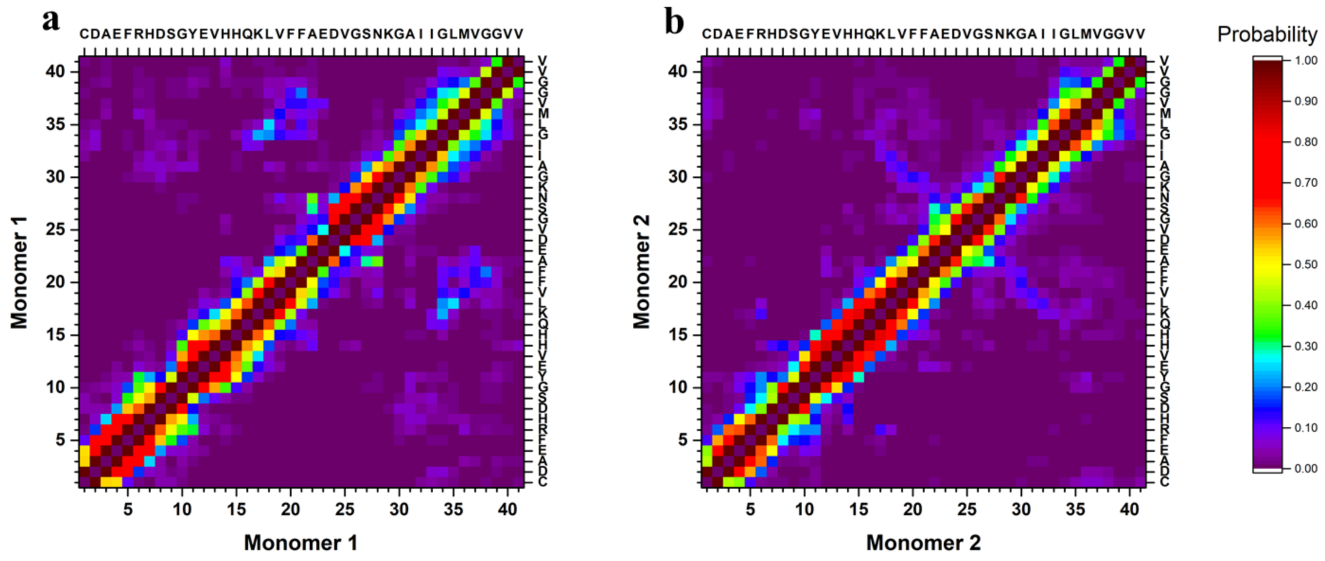


**Figure S4.** Analysis of intra-peptide interactions of Aβ40 monomers within the dimers from 3μs aggregate accelerated MD simulations. Contact probability maps for Cα atoms of Monomer 1, (a), and Monomer 2, (b).

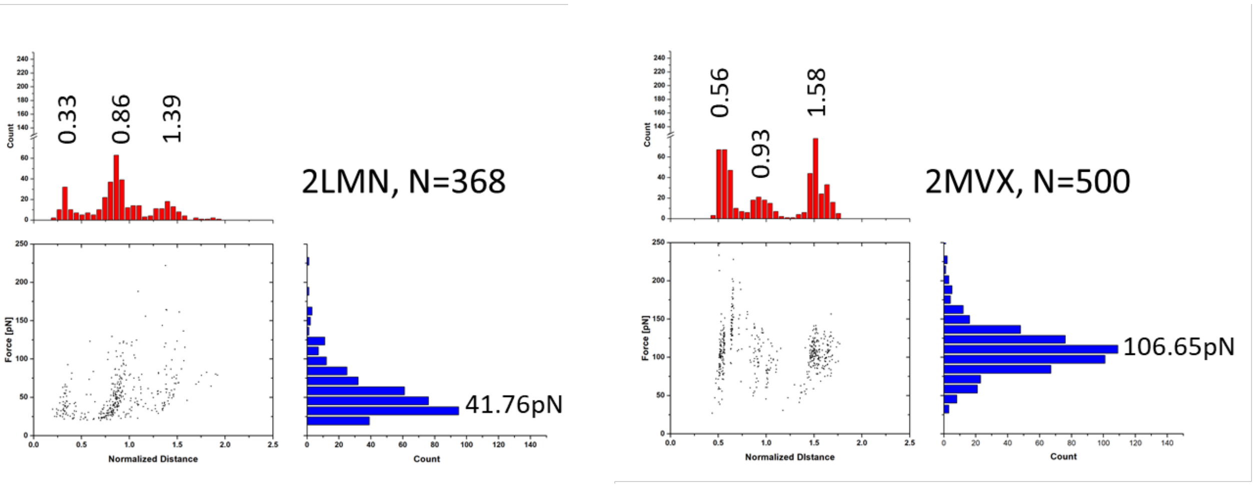


**Figure S5.** Force-induced dissociation results for Aβ40 dimers using MCP simulations. Dimers containing high β-structure content from fibrils (PDB ID: 2LMN and 2MVX). Each dataset shows a scatter plot of Normalized Distance vs Force, a histogram of Force (blue), and a histogram of Normalized Distance (red); normalization was performed based on the experimentally observed contour lengths. Peak values, obtained using Gaussian distribution function, are presented above each peak of the histogram.
